# Evolutionary games of multiplayer cooperation on graphs

**DOI:** 10.1101/038505

**Authors:** Jorge Peña, Bin Wu, Jordi Arranz, Arne Traulsen

## Abstract

There has been much interest in studying evolutionary games in structured populations, often modelled as graphs. However, most analytical results so far have only been obtained for two-player or linear games, while the study of more complex multiplayer games has been usually tackled by computer simulations. Here we investigate evolutionary multiplayer games on graphs updated with a Moran death-Birth process. For cycles, we obtain an exact analytical condition for cooperation to be favored by natural selection, given in terms of the payoffs of the game and a set of structure coefficients. For regular graphs of degree three and larger, we estimate this condition using a combination of pair approximation and diffusion approximation. For a large class of cooperation games, our approximations suggest that graph-structured populations are stronger promoters of cooperation than populations lacking spatial structure. Computer simulations validate our analytical approximations for random regular graphs and cycles, but show systematic differences for graphs with many loops such as lattices. In particular, our simulation results show that these kinds of graphs can even lead to more stringent conditions for the evolution of cooperation than well-mixed populations. Overall, we provide evidence suggesting that the complexity arising from many-player interactions and spatial structure can be captured by pair approximation in the case of random graphs, but that it need to be handled with care for graphs with high clustering.

**Author Summary:** Cooperation can be defined as the act of providing fitness benefits to other individuals, often at a personal cost. When interactions occur mainly with neighbors, assortment of strategies can favor cooperation but local competition can undermine it. Previous research has shown that a single coefficient can capture this trade-off when cooperative interactions take place between two players. More complicated, but also more realistic models of cooperative interactions involving multiple players instead require several such coefficients, making it difficult to assess the effects of population structure. Here, we obtain analytical approximations for the coefficients of multiplayer games in graph-structured populations. Computer simulations show that, for particular instances of multiplayer games, these approximate coefficients predict the condition for cooperation to be promoted in random graphs well, but fail to do so in graphs with more structure, such as lattices. Our work extends and generalizes established results on the evolution of cooperation on graphs, but also highlights the importance of explicitly taking into account higher-order statistical associations in order to assess the evolutionary dynamics of cooperation in spatially structured populations.

## Introduction

Graphs are a natural starting point to assess the role of population structure in the evolution of cooperation. Vertices of the graph represent individuals, while links (edges) define interaction and dispersal neighborhoods. Classical models of population structure, such as island models [1,2] and lattices [3,4], often developed before the current interest in complex networks [5,6], can all be understood as particular instances of graphs [7,8]. More recently, the popularity of network theory has fueled a renewed interest in evolutionary dynamics on graphs, especially in the context of social behaviors such as cooperation and altruism [7–21].

When selection is weak on two competing strategies, such that fitness differences represent only a small perturbation of a neutral evolutionary process, a surprisingly simple condition for one strategy to dominate the other, known as the “sigma rule”, holds for a large variety of graphs and other models of spatially structured populations [22]. Such a condition depends not only on the payoffs of the game describing the social interactions, but also on a number of “structure coefficients”. These coefficients are functions of demographic parameters of the spatial model and of its associated update protocol, but are independent of the payoffs. In the case of two-player games, the sigma rule depends on a single structure coefficient *σ*. The larger this *σ*, the greater the ability of spatial structure to promote the evolution of cooperation or to choose efficient equilibria in coordination games [22]. Partly for this reason, the calculation of structure coefficients for different models of population structure has attracted significant interest during the last years [8,21–27].

Despite the theoretical and empirical importance of two-player games, many social interactions involve the collective action of more than two individuals. Examples range from bacteria producing extracellular compounds [28–31] to human social dilemmas [32–36]. In these situations, the evolution of cooperation is better modeled as a multiplayer game where individuals obtain their payoffs from interactions with more than two players [37–43]. An example of such multiplayer games is the volunteer’s dilemma, where individuals in a group must decide whether to volunteer (at a personal cost) or to ignore, knowing that volunteering from at least one individual is required for a public good to be provided [44–46]. Importantly, such a multiplayer interaction cannot be represented as a collection of pairwise games, because changes in payoff are nonlinear in the number of co-players choosing a particular action.

Multiplayer games such as the volunteer’s dilemma can also be embedded in graphs, assuming, for instance, that nodes represent both individuals playing games and games played by individuals [47–49]. Most previous studies on the effects of graph structure on multiplayer game dynamics have relied on computer simulations [49]. However, similar to the two-player case, some analytical progress can be made if selection is assumed to be weak. In the multiplayer case, the sigma rule depends no longer on one, but on up to *d* − 1 structure coefficients, where *d* is the number of players [50]. Although exact formulas for structure coefficients of multiplayer games can be obtained for relatively simple models such as cycles [51], analysis has proved elusive in more complex population structures, including regular graphs of arbitrary degree. Indeed, extending analytical results on evolutionary two-player games on graphs to more general multiplayer games is an open problem in evolutionary graph theory [52].

Here, we contribute to this body of work by deriving approximate analytical expressions for the structure coefficients of regular graphs updated with a Moran death-Birth model, and hence for the condition of one strategy to dominate another according to the sigma rule. The expressions we find for the structure coefficients suggest that regular graphs updated with a Moran death-Birth model lead to less stringent conditions for the evolution of cooperation than those characteristic of well-mixed populations. Computer simulations suggest that our approximations are good for random regular graphs, but that they systematically overestimate the condition for the evolution of cooperation in graphs with more loops and higher clustering such as rings and lattices. In these cases, cooperation can be no longer promoted, but even be hindered, with respect to the baseline case of a population lacking spatial structure.

## Methods

We consider stochastic evolutionary dynamics on a graph-structured population of size *N*. Each individual is located at the vertex of a regular graph of degree *k*. Individuals obtain a payoff by interacting with their *k* neighbors in a single *d*-person symmetric game (i.e., *d* = *k* + 1). If *j* co-players play *A*, a focal *A*-player obtains *a_j_* whereas a focal *B*-player obtains *b_j_*, as indicated in Table 1.

**Table 1.** Payoffs to *A*-players and *B*-players.

| Opposing $A$-players | 0 | 1 | ... | $j$ | ... | $d-1$ |
| --- | --- | --- | --- | --- | --- | --- |
| payoff to $A$ | $a_0$ | $a_1$ | ... | $a_j$ | ... | $a_{d-1}$ |
| payoff to $B$ | $b_0$ | $b_1$ | ... | $b_j$ | ... | $b_{d-1}$ |

We model the stochastic evolutionary dynamics as a Markov process on a finite space state. More specifically, we consider a Moran death-Birth process [12,14,53] according to which, at each time step: (i) a random individual is chosen to die, and (ii) its neighbors compete to place a copy of themselves in the new empty site with probability proportional to 1 − *w* + *w* × payoff, where the parameter w measures the intensity of selection. Without mutation, such a Markov process has two absorbing states: one where all vertices are occupied by *A*-players and one where all vertices are occupied by *B*-players. Let us denote by *ρ_A_* the fixation probability of a single *A*-player in a population of *B*-players, and by *ρ_B_* the fixation probability of a single *B*-player in a population of *A*-players. We take the comparison of fixation probabilities, i.e.

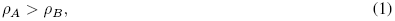

as a measure of evolutionary success [54] and say that *A* is favored over *B* if condition (1) holds.

Under weak selection (i.e., *w* ≪ 1) the condition for *A* to be favored over *B* holds if the sigma rule for multiplayer games [50] is satisfied, i.e., if

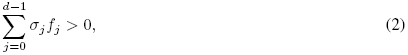

where *σ*_0_,…,*σ*_*d*−1_ are the *d* structure coefficients (constants that depend on the population structure and on the update dynamics), and

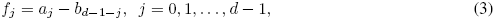

are differences between payoffs, which we will refer to in the following as the “gains from flipping”. The gains from flipping capture the change in payoff experienced by a focal individual playing *B* in a group where *j* co-players play *A* when all players simultaneously switch strategies (so that *A*-players become *B*-players and *B*-players become *A*-players). It turns out that the payoffs of the game only enter into condition (1) via the gains from flipping (3), as the structure coefficients are themselves independent of *a_j_* and *b_j_*.

Structure coefficients are uniquely determined up to a constant factor. Setting one of these coefficients to one thus gives a single structure coefficient for *d* = 2 [22]. For *d* > 2, and in the usual case where structure coefficients are nonnegative, we can impose 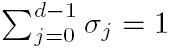 without affecting the selection condition (2). For our purposes, this normalization turns out to be more useful than setting one coefficient to one, as it allows us to rewrite the sigma rule (2) as

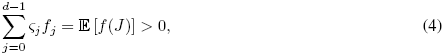

where *f* (*j*) ≡ *f_j_*, and *J* is the random variable with probability distribution prescribed by the “normalized structure coefficients” 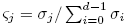. In light of condition (4), the sigma rule can be interpreted as stating that strategy *A* is favored over *B* if the expected gains from flipping are greater than zero when the number of co-players *J* is distributed according to the normalized structure coefficients. From this perspective, different models of population structure lead to different normalized structured coefficients and hence to different expected gains from flipping, which in turn imply different conditions for strategy *A* to be favored over *B* in a given multiplayer game [51]. For instance, a well-mixed population with random group formation updated with either a Moran or a Wright-Fisher process leads to normalized structure coefficients given by [39,40]:

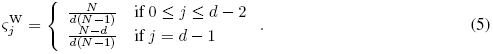

A normalized sigma rule such as the one given by Eq. (4) holds for many spatial models and associated updating protocols [50,51]. Here, we focus on the case of regular graphs updated with a Moran death-Birth process. We provide exact expressions for the case of cycles for which *k* = 2. For *k* ≥ 3, we bypass the difficulties of an exact calculation by using a combination of pair approximation [55,56] and diffusion approximation [14]. Our approach implicitly assumes that graphs are equivalent to Bethe lattices (or Cayley trees) with a very large number of vertices (*N* ≫ *k*). In addition, weak selection intensities (*wk* ≪ 1) are also required for an implicit argument of separation of timescales to hold. In order to assess the validity of our analytical approximations, we implemented a computational model of a Moran death-Birth process in three different types of regular graphs (rings, random graphs, and lattices) with different degrees and estimated numerically the fixation probabilities *ρ_A_* and *ρ_B_* as the proportion of realizations where the mutant succeeded in invading the wild-type.

## Results

### Exact structure coefficients and sigma rule for cycles

Going beyond the complete graph representing a well-mixed population, the simplest case of a regular graph is the cycle, for which *k* = 2 (and consequently *d* = 3). In this case, we find the following exact expressions for the structure coefficients (S1 Text, Section 1):

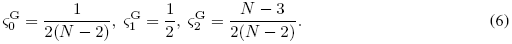

For large *N*, the structure coefficients reduce to 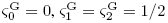 and the sigma rule (4) simplifies to

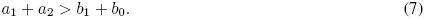

This is also the condition for the boundary between a cluster of *A*-players and a cluster of *B*-players to move in favor of *A*-players for weak selection [57] (Fig. 1). Condition (7) implies that *A* can be favored over *B* even if *A* is strictly dominated by *B* (i.e., *a_j_* < *b_j_* for all *j*) as long as the payoff for mutual cooperation *a*_2_ is large enough so that *a*_2_ > *b*_0_ + (*b*_1_ − *a*_1_); a necessary condition for this inequality to hold is that *A* strictly Pareto dominates *B* (i.e., *a*_2_ > *b*_0_). Such a result is impossible in well-mixed populations, where the structure coefficients (5) prevent strictly dominated strategies from being favored by selection. Condition (7) provides a simple example of how spatial structure can affect evolutionary game dynamics and ultimately favor the evolution of cooperation and altruism.

**Figure 1.**
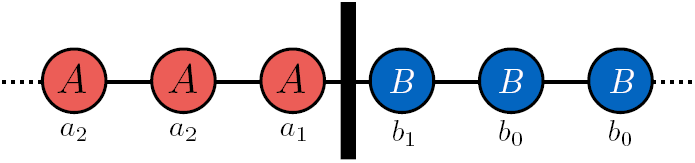
**Payoffs at the boundary of two clusters in the cycle.** Under weak selection, the cluster of *A*-players expands if the sigma rule *a*_1_ + *a*_2_ > *b*_1_ + *b*_0_ holds. As a player is never paired with two players of the opposite strategy, neither *a*_0_ nor *b*_2_ enter into this expression. This provides an intuition behind our analytical results in the simple case when the graph is a cycle.

### **Approximate structure coefficients and sigma rule for regular graphs of degree** *k* ≥ 3

For regular graphs of degree *k* ≥ 3, we find that the structure coefficients can be approximated by (S1 Text, Section 2)

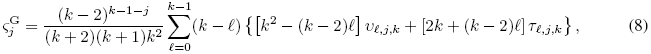

where

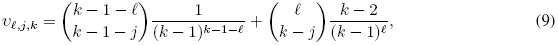

and

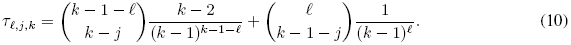

These expressions are nontrivial functions of the degree of the graph *k* and thus difficult to interpret. For instance, for *k* = 3, we obtain 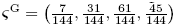.

### Promotion of multiplayer cooperation

The previous results hold for any symmetric multiplayer game with two strategies. To investigate the evolution of multiplayer cooperation, let us label strategy *A* as “cooperate”, strategy *B* as “defect”, and assume that, irrespective of the focal player’s strategy, the payoff of a focal player increases with the number of co-players playing *A*, i.e.,

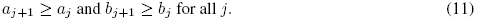

This restriction on the payoffs is characteristic of “cooperation games” [51] in which playing *A* is beneficial to the group but might be costly to the individual. Well-known multiplayer games belonging to this large class of games include different instances of volunteer’s dilemmas [44,46], snowdrift games [58], stag hunts [59], and many other instances of public, club, and charity goods games [43].

We are interested in establishing whether graph-structured populations systematically lead to structure coefficients that make it easier to satisfy the normalized sigma rule (4) than well-mixed populations (the baseline case scenario of a population with no spatial structure) for any cooperation game satisfying condition (11). In other words, we ask whether a graph is a stronger promoter of cooperation than a well-mixed population. Technically, this is equivalent to asking whether the set of games for which cooperation is favored under a graph contains the set of games for which cooperation is favored under a well-mixed population, i.e., whether a graph is greater than a well-mixed population in the “containment order” [51]. A simple sufficient condition for this is that the difference in normalized structure coefficients, ζ^G^ − ζ^W^, has exactly one sign change from − to + [51]. This can be verified for any *N* ≥ 3 in the case of cycles (*k* = 2) by inspection of equations (5) and (6). For large regular graphs of degree *k* ≥ 3 and hence multiplayer games with *d* ≥ 4 players, we checked the condition numerically by comparing equations (5) and (8) for *k* = 3,…, 100. We find that ζ^G^ − ζ^W^ always has a single sign change from − to + and hence that, in the limit of validity of our approximations, regular graphs promote more cooperation than well-mixed populations for all games fulfilling Eq. (11) (Fig. 2). In the following, we explore in more detail the sigma rule for particular examples of multiplayer games.

**Figure 2.**
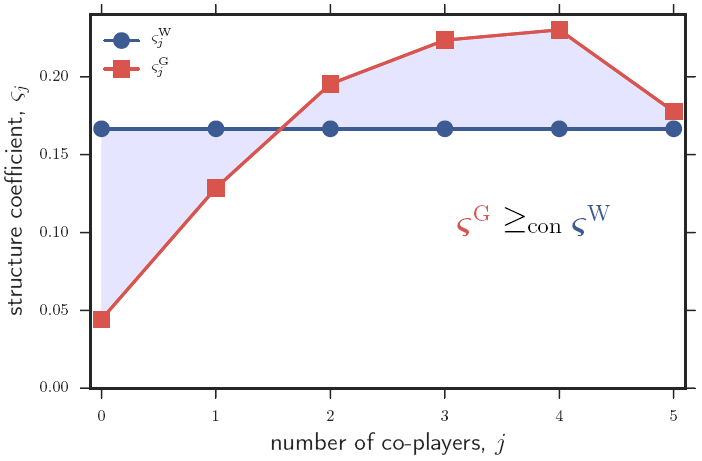
**Structure coefficients and containment order of cooperation.** Approximated (normalized) structure coefficients *ζ^j^* for large regular graphs of degree *k* = 5 updated with a Moran death-Birth process 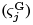 and large well-mixed populations where groups of *d* = 6 players are randomly matched to play a game 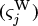. Since *ζ^G^* − *ζ^W^* has one sign crossing from − to +, the graph is greater in the containment order than the well-mixed population (denoted by 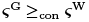). Consequently, if the sigma rule holds for a well-mixed population with coefficients ζ^W^, then it also holds for a graph-structured population with coefficients ζ^G^, for any cooperation game.

### Examples

#### Collections of two-player games

As a consistency check, let us consider the case where individuals play two-player games with their *k* neighbors and collect the payoffs of the different interactions. The two-player game is given by the payoff matrix

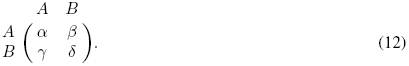

The payoffs for the resulting multiplayer game, which are just the sum of payoffs of the pairwise games, are then given by *a_j_* = *jα* + (*k* − *j*)*β* and *b_j_* = *jγ* + (*k* − *j*)*δ*. The sigma rule (4) can hence be written as

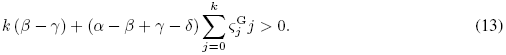

We can show that (S1 Text, Section 2.9)

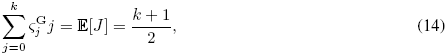

so that condition (13) is equivalent to

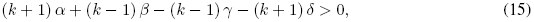

i.e., the sigma rule previously established for pairwise games in regular graphs [cf. Eq. (24) in the Supplementary Material of Ref. [14]]. For a pairwise donation game (for which *α* = *𝓑* − *𝓒*, *β* = *−C*, *γ* = *𝓑*, *δ* = 0, where *𝓑* and *𝓒* are respectively the benefit and cost of donation) this reduces to the well-known *𝓑/𝓒* > *k* rule [7,14,16].

#### Linear games

Suppose now that *a_j_* and *b_j_* are both linear functions of *j*. We can thus write *a_j_* = −*𝓒* + (*𝓑 + 𝓓*) *j/k*, *b_j_* = *𝓑j/k* for some parameters *𝓑*, *𝓒*, and *𝓓*. When *𝓑* > *𝓒* ≥ 0, such a game can be interpreted in terms of a social dilemma as follows. Cooperators each pay a cost *𝓒* in order to provide a benefit *𝓑/k* to each of their co-players; defectors receive the benefits but pay no cost. In addition to the benefit *𝓑/k*, cooperators also get an additional bonus *𝓓/k* per other cooperator in the group. This bonus can be positive or negative.

For such linear games, and by making use of Eq. (14), the sigma condition simplifies to 2*𝓑* + *𝓓*(*k* + 1) > 2*𝓒k*. When there is no bonus (*𝓓* = 0) the game is an additive prisoner’s dilemma [60] and we recover the condition *𝓑/𝓒* > *k*. In the limit of large *k*, the sigma condition becomes *𝓓* > 2*𝓒*.

#### Volunteer’s dilemma

As an example of a nonlinear multiplayer game satisfying condition (11), consider the volunteer’s dilemma [44,45]. In such a game, one cooperator can produce a public good of value *𝓑* at a personal cost *𝓒*; defectors pay no cost and provide no benefit. Payoffs are then given by *a_j_* = *𝓑* − *𝓒* for all *j*, *b*_0_ = 0, and *b_j_* = *𝓑* for *j* > 0. The sigma rule, Eq. (4), for the volunteer’s dilemma hence reduces to

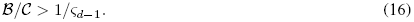

For the cycle, we thus find

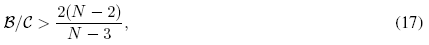

which in the limit of large *N* reduces to *𝓑/𝓒* > 2. For large regular graphs of degree *k* ≥ 3, our approximations lead to

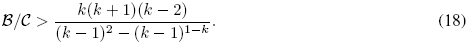

These conditions contrast with that for a large well mixed population, which is given by *𝓑/𝓒* > *k* + 1.

Suppose now that the cost of producing the public good is shared among cooperators [46]. Payoffs are then given by *a_j_* = *𝓑* − *𝓒*/(*j* + 1), *b*_0_ = 0 and *b_j_* = *𝓑* for *j* > 0. In this case the sigma rule simplifies to

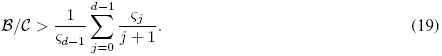

This leads to

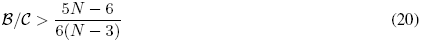

in the case of a finite cycle of size *N* and *𝓑/𝓒* > 5/6 for a large cycle. Contrastingly,in a well-mixed population,

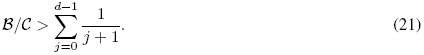

### Computer simulations

To assess the validity of our approximations, we compare our analytical results with explicit simulations of evolutionary dynamics on graphs (Fig. 3, *N* = 100; S1 Fig, *N* = 500). We implemented three different kinds of regular graphs: (i) random regular graphs, (ii) rings (generalized cycles in which each node is connected to *k*/2 nodes to the left and *k*/2 nodes to the right), and (iii) lattices (a square lattice with von Neumann neighborhood with *k* = 4, a hexagonal lattice with *k* = 6, and a square lattice with Moore neighborhood and *k* = 8). Analytical predictions are in good agreement with simulation results in the case of cycles (i.e., rings with *k* = 2, for which our expressions are exact) and for all random regular graphs that we explored. Contrastingly, for rings with *k* ≥ 4 and lattices, our approximations tend to underestimate the critical benefit-to-cost ratio beyond which the fixation probability of cooperators is greater than that of defectors. In other words, our analytical results seem to provide necessary but not sufficient conditions for cooperation to be favored. Such discrepancies stem from the fact that our analysis assumes graphs with no loops such as Cayley trees; the error induced by our approximations is more evident when looking at the actual fixation probabilities (S2 Fig, *N* = 100, S3 Fig, *N* = 500) and not just at their difference. As all graphs with *k* > 2 we considered do contain loops, such mismatch is expected—in particular for rings and lattices, which are characterized by high clustering. Perhaps more importantly, our simulations suggest that the critical benefit-to-cost ratio for the volunteer’s dilemma without cost sharing in rings and lattices with *k* ≥ 6 is greater than the corresponding values for random graphs and well-mixed populations. This illustrates a case in which a graph-structured population updated with a death-Birth process leads to less favorable conditions for the evolution of cooperation than a well-mixed population.

## Discussion

We studied evolutionary multiplayer game dynamics on graphs, focusing on the case of a Moran death-Birth process on regular structures. First, we used a combination of pair approximation and diffusion approximation to provide analytical formulas for the structure coefficients of a regular graph, which together with the payoffs from the game determine when a strategy is more abundant than another in the limits of weak selection and weak mutation. Such a condition is valid for any symmetric multiplayer game, including the volunteer’s dilemma [44–46] and other multiplayer social dilemmas discussed in the recent literature [38,41,58,59,61]. The condition can be used to determine the specific conditions (in terms of the degree of the graph and the parameters of the game, such as payoff costs and benefits) under which cooperation will thrive. The structure coefficients also provide a way of comparing the graph with other population structures, such as the well-mixed population. In particular, and to the extent that our approximations are valid, graphs updated with a death-Birth process are more conducive to the evolution of cooperation than well-mixed populations for a large class of games (see condition (11)).

**Figure 3.**
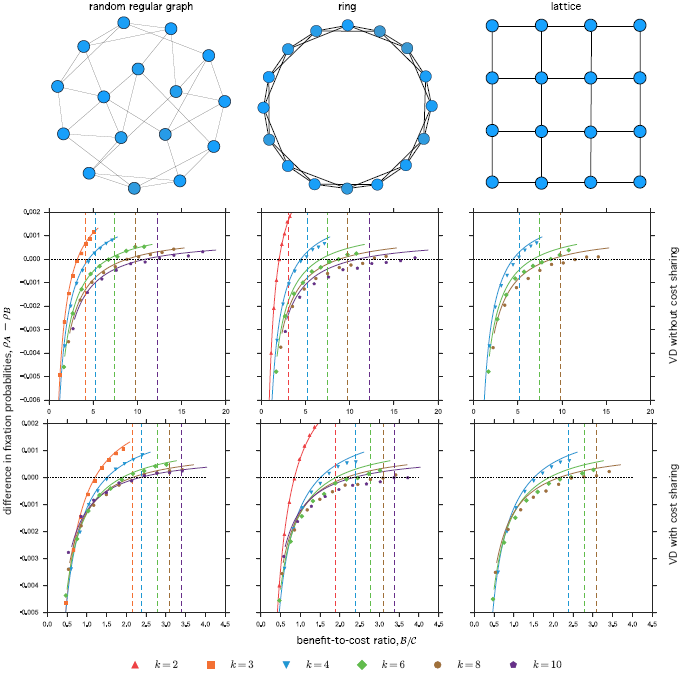
**Simulations of evolutionary game dynamics on graphs (difference in fixation probabilities,** *N* = 100). The first row shows the type of (regular) graph for the particular case of *k* = 4, i.e., each node has exactly four neighbors. The second and third rows show simulation results for the volunteer’s dilemma without cost-sharing and with cost-sharing, respectively. Simulation data in the first column correspond to random regular graphs, in the second column to rings, and in the third column to lattices. The fixation probability of cooperators, *ρ_A_* (defectors, *ρ_B_*) was calculated as the fraction of runs where a single cooperator (defector) reached fixation out of 10^7^ runs. Symbols show the difference between such fixation probabilities, as a function of the benefit-to-cost ratio *𝓑/𝓒*, for different types and degrees of the graph. Lines indicate analytical predictions for the difference in fixation probabilities (left hand side of Eq. (4) with normalized sigmas given by Eq. (6) or Eq. (8)). Dashed vertical lines the critical benefit-to-cost ratios *𝓑/𝓒* above which we have *ρ_A_* > *ρ_B_* for well-mixed populations (right hand side of Eq. (16) or Eq. (19) with normalized sigmas given by Eq. (5)). Parameters: population size *N* = 100, intensity of selection *w* = 0.01, payoff cost *𝓒* = 1.

Second, we used numerical simulations to estimate the fixation probabilities and the difference in fixation probabilities of different strategies for particular examples of games (volunteer’s dilemma with and without cost sharing) and graphs (random regular graphs, rings, and lattices). Although simulations agree very well with the analytical approximations in the case of random regular graphs, discrepancies are evident in the case of rings and lattices, which are characterized by higher clustering and for which pair approximation is not sufficiently accurate. In these cases, the analytical approximations systematically overestimate the ability of a graph to promote the evolution of cooperation. Importantly, in the case of the volunteer’s dilemma without cost sharing and for rings or lattices of relatively large degree, the critical benefit-to-cost ratio above which cooperation is favored is greater, not smaller, than the corresponding value for a well-mixed population. Even though detrimental effects of spatial structure on cooperation have been previously noted in similar studies [62], our results are counterintuitive given the updating protocol and the intensity of selection we explored. Indeed, a death-Birth Moran process under weak selection would always favor cooperation (with respect to a well-mixed population of the same size) for any linear cooperation game, including any collection of two-player cooperation games. Our simulations show that this might not be the case when social dilemmas are instead modelled as nonlinear games such as the volunteer’s dilemma.

We used pair approximation and diffusion approximation to find approximate values for the structure coefficients, but other approaches can be used to obtain better estimates of them. In particular, coalescent theory [63] allows us to write the sigma rule in terms of selection coefficients (dependent on the payoffs of the game and the demographic parameters of the model) and expected coalescence times under neutrality [64,65]; however, such expected coalescence times can be difficult to obtain exactly. Alternatively, for small graphs, the sigma rule and hence the structure coefficients can be explicitly calculated from the transition matrix of the evolutionary process (cf. Appendix C of Ref. [26]). Finally, we note that even in cases for which the structure coefficients are difficult to obtain by purely analytical means, they can be estimated numerically, either indirectly (by estimating the expected times to coalescence) or directly (by computing and comparing fixation probabilities).

For simplicity, we assumed that a focal player obtains its payoff from a single multiplayer game with its *k* immediate neighbors. Such assumption allowed us to consider multiplayer interactions on graphs in a straightforward way. However, this is in contrast with a common assumption of many studies of multiplayer spatial and network games in which a focal player’s total payoff is the sum of payoffs obtained in *k* + 1 different games, one “centered” on the focal player itself and the other *k* centered on its neighbors [47–49]. As a result, focal players interact not only with first-order but also with second-order neighbors, which would lead to more intricate structure coefficients. For example, in this case the structure coefficients of a cycle are given by [51,66]

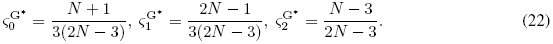

These values are different from those we calculated under the assumption that individuals play a single game with first-order neighbors, given by Eq. (6). For *N* > 4, the structure coefficients fulfill 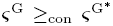, meaning that our assumption of payoffs from a single game leads to less restrictive conditions for cooperation to be favored by selection. This observation is in line with previous results for pairwise games on graphs suggesting that the condition for the evolution of cooperation is optimized when interaction and replacement neighborhoods coincide [67], which corresponds to our assumption of individuals playing a single game. Future work should consider the calculation of structure coefficients for the cases where the payoff to a player also depends on games centered on neighbors and how the condition for the promotion of cooperation differs from the one resulting from our simplifying assumption.

We modelled social interactions as multiplayer matrix games with two discrete strategies (*A* and *B*) and obtained our results by assuming that selection is weak (*w* is small). Alternatively, one could model the same multiplayer game but assume instead that players can choose between two similar mixed strategies *z* and *z* + *δ*, where *z* and *z* + *δ* refer to the probability of playing *A* for each strategy, and *δ* is small [43,68,69]. In such a “*δ*-weak selection” scenario, and for any number of players, only a single structure coefficient is needed to identify conditions under which a higher probability of playing *A* is favored by natural selection. For transitive graphs of size *N* and degree *k*, this structure coefficient is given by [7,25]

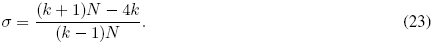

Exchanging the structure coefficient *σ* for the “scaled relatedness coefficient” *κ* of inclusive fitness theory via the identity *κ* = (*σ* − 1)/(*σ* + 1) [65], we obtain [16]

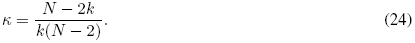

With such a value, recent results on multiplayer discrete games in structured populations under *δ*-weak selection [43] can be readily applied to show that, for all cooperation games as we defined them and for a death-Birth protocol, *A* is favored over *B* more easily for a graph-structured population than for a well-mixed population, as long as *N* > *k* + 1. Such prediction qualitatively coincides with the one obtained from our analytical approximations, but does not capture our numerical results for the volunteer’s dilemma in rings and lattices.

To sum up, we have shown that even for multiplayer games on graphs, which are routinely analyzed by simulation only, some analytical insight can be generated. However, fully accounting for the complexity of evolutionary multiplayer games in graphs with high clustering remains a challenging open problem.

## Supporting Information

### S1 Text

#### Supplementary Methods

Calculations for the exact structure coefficients of cycles, the approximate structure coefficients of graphs of degree *k* ≥ 3, and a short description of the computational model used for our simulations.

### S1 Fig

**Simulations of evolutionary game dynamics on graphs (difference in fixation probabilities,** *N* = 500). Same as in Fig. 3, but for a population size *N* = 500.

### S2 Fig

**Simulations of evolutionary game dynamics on graphs (fixation probabilities,** *N* = 100). Open symbols show the fixation probability of cooperators (*ρ_A_*) and filled symbols the fixation probability of defectors (*ρ_B_*) as a function of the benefit-to-cost ratio *𝓑/𝓒*, for different types and degrees of the graph. Lines indicate analytical predictions for the fixation probabilities. Parameters: population size *N* = 100, intensity of selection *w* = 0.01, payoff cost *𝓒* = 1.

### S3 Fig

**Simulations of evolutionary game dynamics on graphs (fixation probabilities,** *N* = 500). Same as in S2 Fig, but for a population size *N* = 500.

## Supplementary Methods

### 1 The cycle

We start by considering the cycle, for which *k* = 2, and where a single mutant always leads to a connected cluster of mutants. In this case, analytical expressions for the fixation probabilities of the two types and for the structure coefficients can be obtained exactly by adapting previous results on two-player games on cycles [1]. The state space of the stochastic process is *i* = 0,…, *N*, where *i* is the number of *A*-players and *N* is the population size. At each time step, the number of *i* players either increases by one (with probability 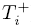), decreases by one (with probability 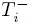), or remains the same (with probability 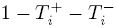). The fixation probability of a single mutant *A* is given by [2,3]

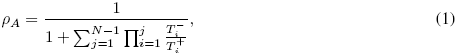

and the ratio of the fixation probabilities is given by [2,3]

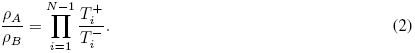

In order to compute these quantities, we need to find expressions for the ratio of the transition probabilities, 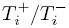, for each *i* = 1,…, *N* − 1. For convenience, let us define *α_j_* = 1 − *w* + *wa_j_* and *β_j_* = 1 − *w* + *wb_j_* for *j* = 0,1,2, where *w* is the intensity of selection and *a_j_* (*b_j_*) is the payoff of an *A*-player (*B*-player) when playing against two other players, *j* ∈ {0,1,2} of which are *A*-players. For a death-Birth protocol, we find

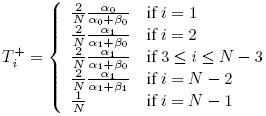

and

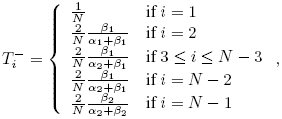

so that the ratio of transition probabilities is given by

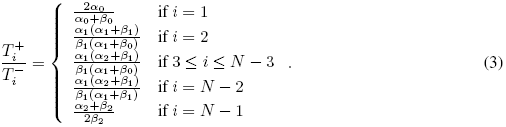

With the previous expression, and for weak selection (*w* ≪ 1) we obtain

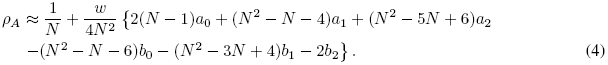

By symmetry, the expression for *ρ_B_* can be obtained from the expression for *ρ_A_* after replacing *a_j_* by *b_k−j_* and *b_j_* by *a_k−j_*, i.e.,

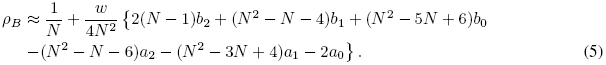

In a similar manner, for weak selection the ratio of fixation probabilities can be approximated by

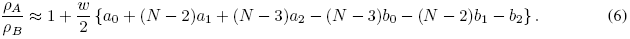

Thus, the condition *ρ_A_* > *ρ_B_* becomes

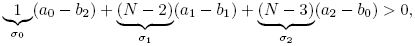

from which we identify the structure coefficients:

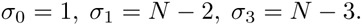

As we assume *N* ≥ 3, the structure coefficients are nonnegative. Normalizing the structure coefficients we obtain

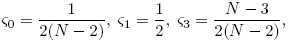

which are the values given by Eq. (6) in the main text.

### 2 Regular graphs with *k* ≥ 3

For regular graphs with degree *k* ≥ 3, we obtain the structure coefficients by finding an approximate expression for the comparison of fixation probabilities, *ρ_A_* > *ρ_B_*. To estimate these fixation probabilities, we follow closely the procedure used by Ohtsuki et al. [4], based on a combination of pair approximation and diffusion approximation.

#### 2.1 Pair approximation

Let us denote by *p_A_* and *p_B_* the global frequencies of types *A* and *B*, by *p_AA_*, *p_AB_*, *p_BA_* and *p_BB_* the frequencies of *AA*, *AB*, *BA*, and *BB* pairs, and by *q_X|Y_* the conditional probability of finding an *X*-player given that the adjacent node is occupied by a *Y*-player, where *X* and *Y* stand for *A* or *B*. Such probabilities satisfy

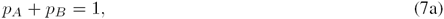

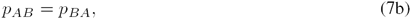

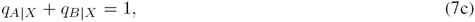

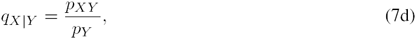

implying that the system can be described by only two variables: *p_A_* and *p_AA_*.

In probabilistic cellular automata such as the one analyzed here, the dynamics of frequencies of types (*p_A_*, *p_B_*) and pairs of types (*p_AA_*, *p_AB_*, *p_BA_* and *p_BB_*) will depend on triplets and higher-order spatial configurations [5]. Pair approximation allows us to obtain a closed system by approximating third- and higher-order spatial moments by heuristic expressions involving second- and first-order moments only [5,6]. In particular, for each site *X*, we assume that the probability of finding *j A*-players among its *k* neighbors follows the binomial distribution

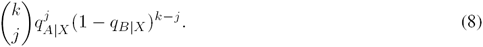

Likewise, for each pair *XY*, we assume that the probability of finding *j A*-players among the *k* − 1 neighbors of *X* not including *Y* follows the binomial distribution

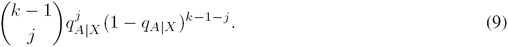

Here, we implicitly assume that members of a pair are unlikely to have common neighbors, as it is approximately the case for random graphs. In this case, Eq. (8) and Eq. (9) are standard (and parsimonious) assumptions (cf. Ref. [5], Eq. 19. 27). For other graphs (such as lattices) the overlap among the neighbors of a pair introduce correlations not taken into account by our simplification.

In the following, we write down the change of *p_A_* and *p_AA_* under the assumptions of pair approximation. Then, we assume that selection is weak and that a separation of timescales holds in order to reduce the dimension of the system of equations. Finally, we employ a diffusion approximation to get the equation that governs the fixation probabilities. From the expressions of the fixation probabilities, the structure coefficients can be obtained after some cumbersome algebra.

#### 2.2 Updating a *B*-player

A *B*-player is chosen to die with probability *p_B_*; its *k* neighbors compete for the vacant vertex proportionally to their effective payoffs. Denoting by *k_A_* and *k_B_* the number of *A* and *B* players among these *k* neighbors, and by virtue of Eq. (8), the frequency of such configuration is given by

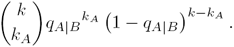

The effective payoff of each *A*-player connected by an edge to the dead *B*-player is given by

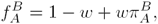

where

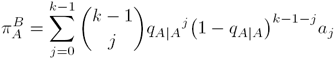

is, by virtue of Eq. (9), the expected payoff to an *A*-player with one *B* co-player and *k* − 1 other players.

Likewise, the effective payoff of each *B*-player connected by an edge to the dead *B*-player is given by

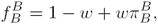

where

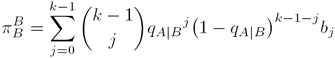

is the expected payoff to a *B*-player with one *B* co-player and *k* − 1 other players.

Under weak selection, the probability that a neighbor playing *A* replaces the vacant spot left by the dead *B*-player is given by

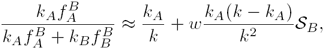

where

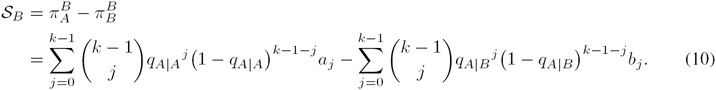

Hence, the frequency *p_A_* of *A*-players in the population increases by 1/*N* with probability

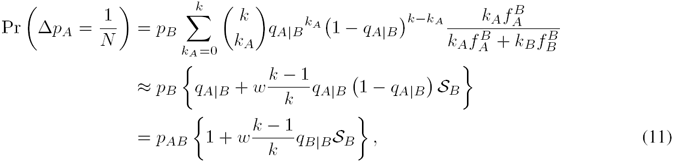

where we used the formulas for the first two moments of a binomial distribution and the identities *p_B_q_A|B_* = *p_AB_* and 1 − *q_A|B_* = *q_B|B_* implied by Eq. (7).

Regarding pairs, if the *B*-player chosen to die is replaced by an *A*-player then the number of *AA* pairs increases by *k_A_*. Since the total number of pairs in the population is equal to *kN*/2, the proportion *p_AA_* of *AA* pairs increases by 2*k_A_*/(*kN*) with probability

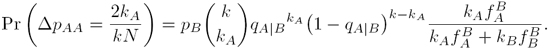

#### 2.3 Updating an *A*-player

An *A*-player is chosen to die with probability *p_A_*. There are *k_A_ A*-players and *k_B_ B*-players in the neighborhood of the vacant node. The frequency of this configuration is (cf. Eq. (8))

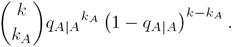

The effective payoff of each neighboring *A*-player is

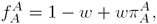

where

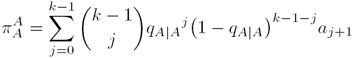

is, by virtue of Eq. (9), the expected payoff to an *A*-player with one *A* co-player and *k* − 1 other players.

Likewise, the effective payoff to each neighboring *B*-player is given by

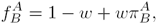

where

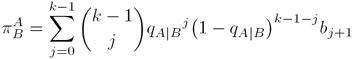

is the expected payoff to a *B*-player with one *A* co-player and *k* − 1 other players.

The probability that one of the neighbors playing *B* replaces the vacancy is given by

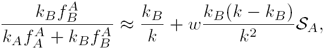

where

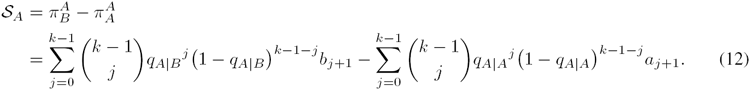

The vacancy is replaced by a *B*-player and therefore *p_A_* decreases by 1/*N* with probability

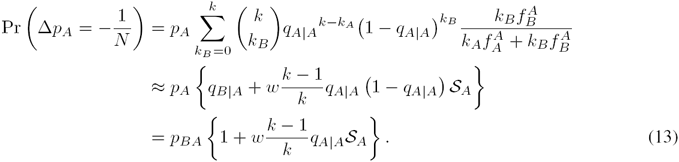

Regarding pairs, the proportion *p_AA_* of *AA* pairs decreases by 2*k_A_*/(*kN*) with probability

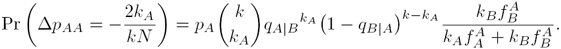

#### 2.4 Separation of time scales

Supposing that one replacement event takes place in one unit of time, the time derivative of *p_A_* is given by

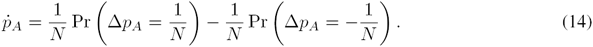

Using Eq. (11) and (13) we obtain, to first order in *w*:

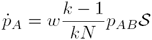

where

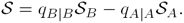

Similarly to Eq. (14), the time derivative of *p_AA_* is given by

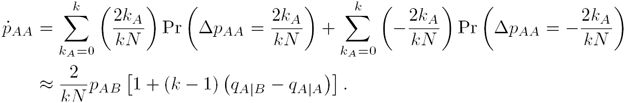

For weak selection (*wk* ≪ 1) the local density *p_AA_* equilibrates much more quickly than the global density *p_A_*. Therefore, the dynamical system rapidly converges onto the slow manifold where *ṗ_AA_* = 0 and hence

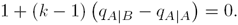

From this expression and Eq. (7) we obtain

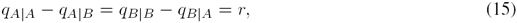

where we define

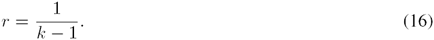

As pointed out by Ohtsuki et al. [4], Eq. (15) measures the amount of positive correlation or effective assortment between adjacent players generated by the population structure. Moreover, expression (15) together with Eq. (7) leads to

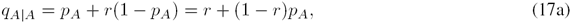

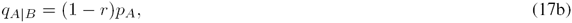

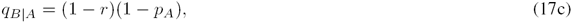

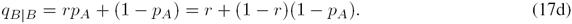

#### 2.5 Algebraic manipulations

It follows from the previous approximations that *ṗ_A_* is proportional to

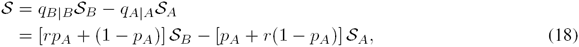

which is a polynomial of degree *k* in *p_A_*. Let us write such polynomial in a more compact form. To do so, we make use of the following identities:

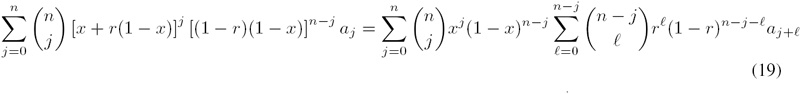

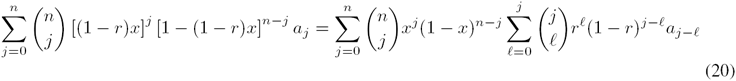

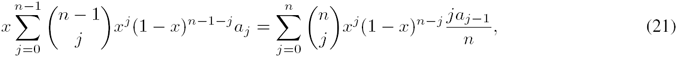

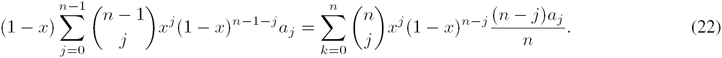

Proofs of identities (21) and (22) are provided in Appendix B of Ref. [7]. In the following, we prove (19) [(20) is proven in a similar way]. Starting from the left side of (19) we expand the term [*x* + *r*(1 − *x*)]*^j^* (by applying the binomial theorem) and rearrange to obtain:

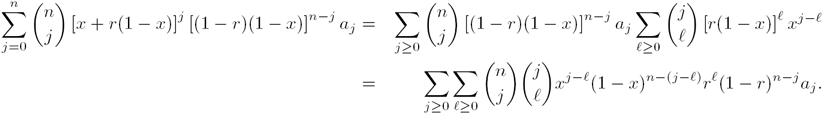

Now, since

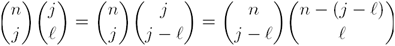

and introducing *m* = *j* − *ℓ*, we can write

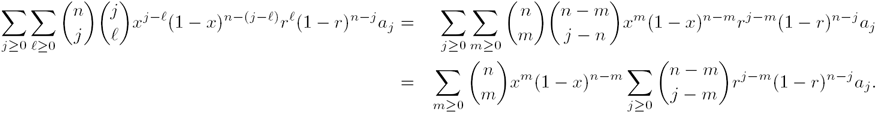

Replacing *ℓ* = *j* − *m* in the last sum,

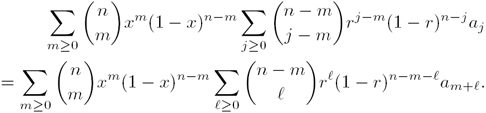

Finally, changing the dummy variable *m* by *j* in the last expression and making explicit the upper limits of the sums, we obtain the right side of (19).

Replacing Eq. (17a) and (17b) into Eq. (10), and applying identities (19) and (20), we can write

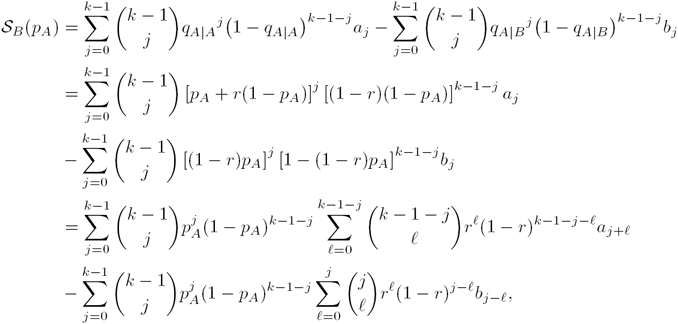

so that we obtain

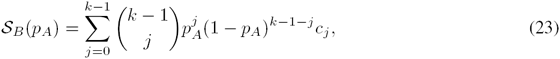

where

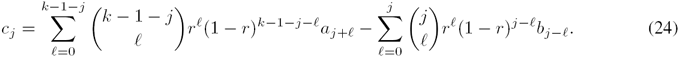

Likewise, replacing Eq. (17) into Eq. (12), and applying identities (19) and (20) we obtain

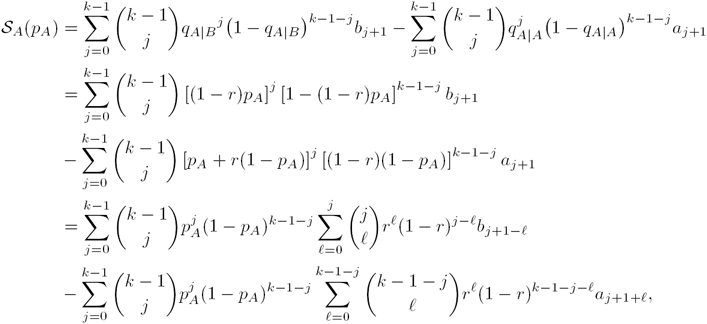

and hence

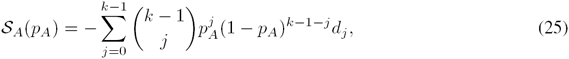

where

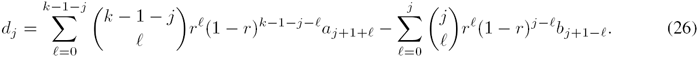

Replacing Eq. (23) and Eq. (25) into Eq. (18), and applying identities (21) and (22), we finally obtain

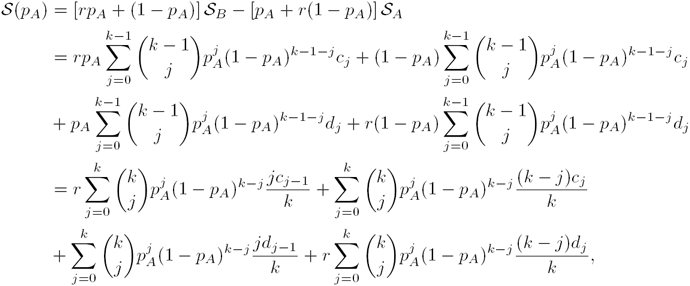

and hence

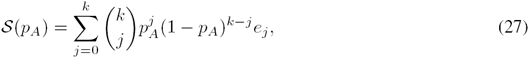

where

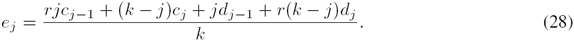

#### 2.6 Diffusion approximation

Assuming that Eq. (17) holds, we study a one dimensional diffusion process on the variable *p_A_*. Therefore, within a short interval, Δ*t*, we have

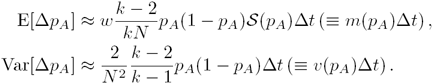

The fixation probability, *ϕ_A_*(*y*) of strategy *A* with initial frequency *p_A_* (*t* = 0) = *y*, is then governed by the differential equation (cf. Eq. 4.13 in Ref. [8])

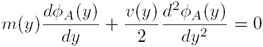

with boundary conditions *ϕ_A_*(0) = 0 and *ϕ_A_*(1) = 1.

The probability that absorption eventually occurs at *p_A_* = 1 is then (cf. Eq. 4.17 in Ref. [8])

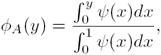

where (cf. Eq. 4.16 in Ref. [8])

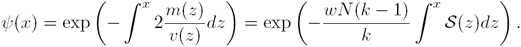

Since we assume that *w* is very small,

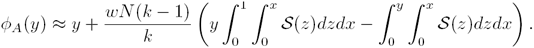

This expression involves integrals of *𝓢*(*z*). Using the formula for the integral of a polynomial in Bernstein form (cf. p. 391 of Ref. [9]), i.e.,

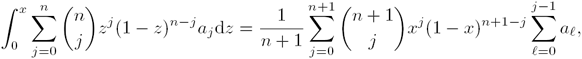

we obtain

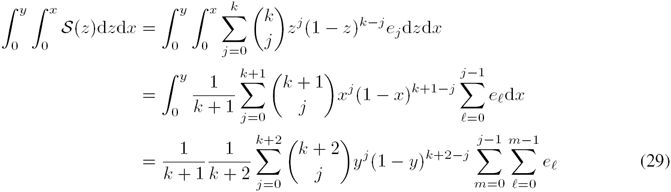

and hence

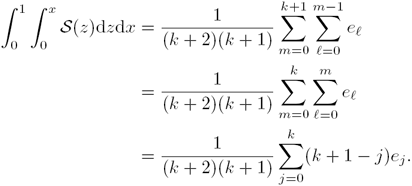

Writing out Eq. (29) as

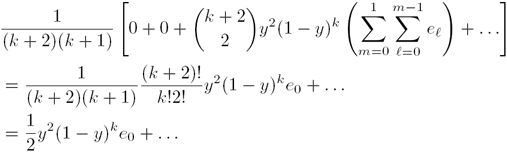

it is clear that, for *y* = 1/*N* and *N* large, 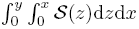 can be approximated by

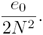

The fixation probability, *ρ_A_* = *ϕ_A_*(1/*N*), can then be written as

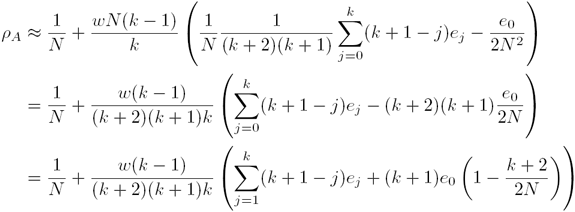

If *k* ≪ *N*, then (*k* + 2)/(2*N*) ≪ 1, and we finally obtain

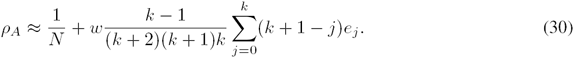

#### 2.7 Fixation probabilities, sigma rule and structure coefficients

From Eq. (30), the fixation probability of a mutant *A* is greater than neutral if 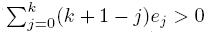. By Eq. (28), the coefficients *e_j_* are linear in *c_j_* and *d_j_*, which are linear in the payoff entries *a_j_* and *b_j_* (cf. Eq. (24) and (26)). Thus, 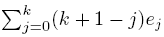 is linear in the payoff entries, meaning that there exist *α_j_* and *β_j_* such that

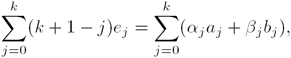

and so

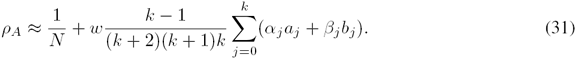

By symmetry, the fixation probability of a single *B* mutant is given by

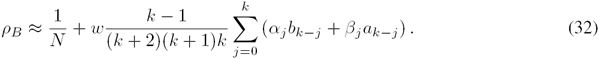

Therefore, under weak selection

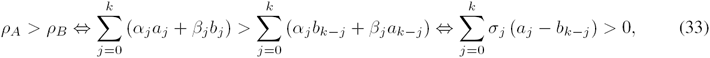

where

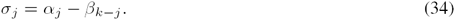

The rightmost expression in Eq. (33) has been termed the “sigma rule” and the coefficients *σ_j_* are the structure coefficients [10–12].

To obtain expressions for the fixation probabilities and the structure coefficients, we need to calculate *α_j_* and *β_j_*. For a multiplayer game with *a_j_* = *δ_i,j_* (i.e., *a_j_* = 1 for some *j* = *i* and *a_j_* = 0 otherwise), we have

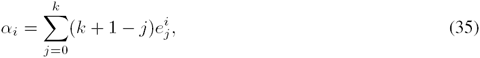

where 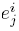 denotes the coefficient *e_j_* with *a_j_* = *δ_i,j_* and *b_j_* = 0 for all *j*. Replacing the formula for *e_j_* (28) into Eq. (35), expressing *r* in terms of *k* (Eq. (16)), and simplifying we get

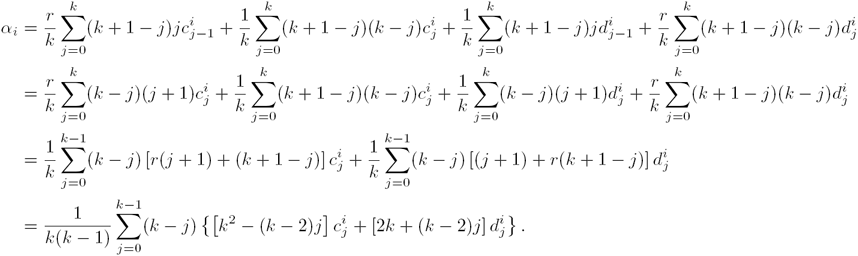

Now, since for *α_i_* we have that *a_j_* = *δ_i,j_* and *b_j_* = 0 for all *j*, and from Eq. (24), (26), and Eq. (16), we have

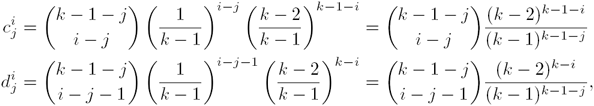

and 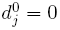 for all 0 ≤ *j* ≤ *k* − 1. Hence

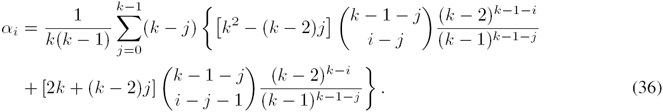

Similarly (now letting the payoff entries be *b_j_* = *δ_i,j_* and *a_j_* = 0) we obtain

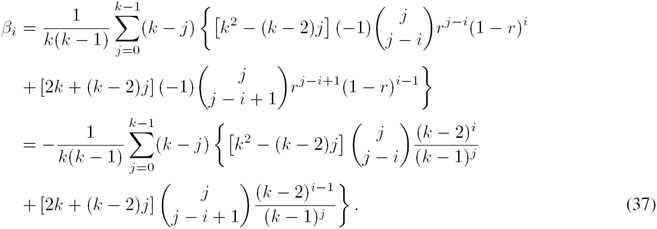

Replacing expressions (36) and (37) into Eq. (34) and simplifying, we finally obtain the following expressions for the structure coefficients

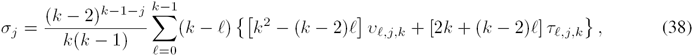

where

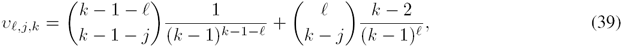

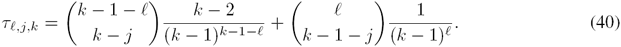

#### 2.8 Normalized structure coefficients

The structure coefficients given by Eq. (38) are nonnegative. Once we have an expression for their sum, we can normalize the structure coefficients so that they describe a probability distribution. In the following we work out such an expression.

We start by noting that, subtracting Eq. (32) from Eq. (31), the difference of the fixation probabilities under our approximations can be written as

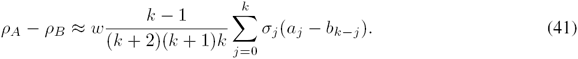

In particular, this expression holds for a multiplayer game with payoffs given by *a_j_* = 1 and *b_j_* = 0 for all *j*, for which Eq. (41) reduces to

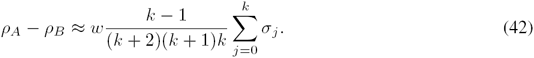

The multiplayer game with payoffs *a_j_* = 1 and *b_j_* = 0 for all *j* is mathematically equivalent to a collection of pairwise games played with neighbors with a payoff matrix

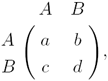

where *a* = *b* = 1/*k* and *c* = *d* = 0. Indeed, for such payoff values the accumulated payoff to an *A*-player is always 1 and that of a *B*-player is always 0. For a general pairwise game, we have that (cf. Eqs. (19) and (21) in the Supplementary Material of Ref. [4])

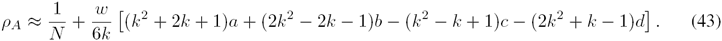

By symmetry:

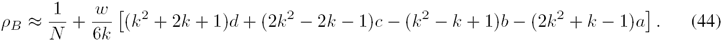

Therefore, for *a* = *b* = 1/*k* and *c* = *d* = 0, we have that

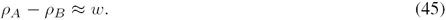

Since the right hand side of Eq. (42) should be equal to the right hand side of Eq. (45), we conclude that

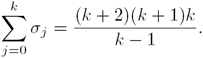

Defining

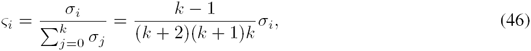

we finally obtain

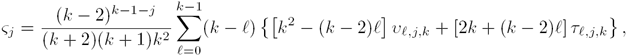

which is the expression for the normalized structure coefficients as given in Eq. (8) of the main text.

#### 2.9 A useful identity

If individuals play the pairwise game

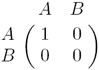

with each neighbor, then by Eqs. (43) and (44) we have

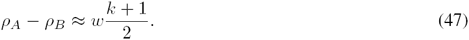

Now consider the difference in fixation probabilities arising from the equivalent multiplayer version, for which *a_j_* = *j* and *b_j_* = 0 for all *j*. Replacing *a_j_* = *j* and *b_j_* = 0 into Eq. (41) leads to

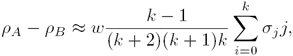

which by Eq. (46) can be written as

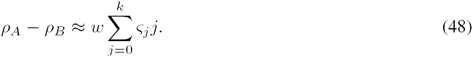

in terms of the normalized structure coefficients. Comparing Eq. (47) and (48), we finally obtain

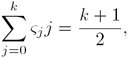

which is the expression in Eq. (14) in the main text. Note that this expression is valid only in the limit of large *N*.

### 3 Computional model

We implemented numerical simulations of a Moran process with death-Birth (dB) updates for different kinds of graphs. The simulations rely on three different types of graphs: random regular, ring and lattice. We employ the C version of the igraph library^1^ to generate all random regular graphs —igraph_k_regular_game()—, the ring of degree *k* = 2 —igraph_ring()— and the lattice of degree *k* = 4 —igraph_lattice(). Given that igraph does not provide generators for lattices of *k* > 4, we implemented an algortihm that extends a lattice of degree *k* = 4 (von Neumann neighborhood) to degrees *k* = 6 (hexagonal lattice) and *k* = 8 (Moore neighborhood). Similarly, we extend the ring of degree *k* = 2 by increasing its connectivity accordingly to generate cycles of degrees *k* = 4, 6, 8, 10.

At each realization of the simulation we start with a monomorphic population playing one of the two strategies and add a single mutant of the opposite strategy in a randomly chosen vertex. We allow the simulation to run until it reaches an absorbing state (i.e., when either of the two strategies reaches fixation). At each simulation step a vertex (*a*) is randomly selected from the whole population (i.e., death step) and a second vertex (*b*) is selected from the neighborhood of *a* with a probability proportional to its fitness. During this step we use the stochastic acceptance algorithm [13] to select an individual with a probability proportional to its fitness. Hereafter, the strategy of vertex *b* is copied to *a* (i.e., death step). The payoffs of the nodes —which depend on the game in place, their own strategies, and the strategies of their neighbours— are calculated as discussed in the main text. For optimization purposes, in the first step of each realisation we compute the payoffs of the whole network. Thenceforth we only re-compute the payoff of a vertex and its neighbors whenever a vertex switches its strategy.

We repeat this process for 10^7^ different realizations and keep track of the number of times the mutant strategy has reached fixation. At the final step we compute the fixation probability of the mutant strategy as the ratio between the number of hits —i.e., number of times the mutant invaded the resident strategy— and the total number of realizations. We run separate simulation batches for both strategies, in a way that both strategies play as the mutant and resident.

## Footnotes

1 http://igraph.org/

